# A life-history perspective on the evolutionary interplay of sex ratios and parental sex roles

**DOI:** 10.1101/2022.11.23.517539

**Authors:** Xiaoyan Long, Tamas Székely, Jan Komdeur, Franz J. Weissing

## Abstract

Parental care is one of the most diverse social behaviours, and caring by the male, female or both parents is essential for successful reproduction of many organisms. Theoretical and empirical studies suggest that parental sex roles are associated with biased sex ratios. However, there is considerable debate on the causal relationship between parental sex roles and sex ratio biases and on the relative importance of the operational sex ratio (OSR), the adult sex ratio (ASR), and the maturation sex ratio (MSR). Here we use individual-based evolutionary simulations to investigate the joint evolution of sex-specific parental behaviour and the various sex ratios in several life history scenarios. We show that sex differences in maturation time or mortality rates at various life-history stages predict the evolution of parental sex roles relatively well: typically, but not always, parental care is biased toward the sex with the lower mortality or the faster maturation. The association of parental sex roles with the various sex ratios is more intricate. In our simulations, the operational sex ratio at evolutionary equilibrium was typically biased toward the less-caring sex. However, the direction and strength of OSR biases often changed drastically in the course of evolution, implying that, rather than being a driver of parental sex roles, OSR biases emerge as a consequence of sex-biases in parental care. When the MSR or the ASR is biased, this bias is generally associated with a bias in parental care: the overrepresented sex does most of the caring. However, the opposite pattern (that the underrepresented sex did most of the caring) also occurs in some scenarios. Moreover, pronounced parental sex roles may also evolve in the absence of an MSR or ASR bias. Taken together, we conclude that none of the sex ratios can be viewed as drivers of a parental care bias; they rather co-evolve with parental care bias in a subtle manner.

## Introduction

When animal species provide post-zygotic parental care, the degree to which each sex contributes varies strongly across taxa (Clutton-Brock, 1991; Balshine, 2012; Trumbo, 2012). Females provide most of the care in mammals and invertebrates (Balshine, 2012; Trumbo, 2012), both sexes are involved in parental duties in the majority of avian species (Cockburn, 2006; Balshine, 2012), and male-biased or male-only care is common in fishes with parental care (Blumer, 1979; Mank et al., 2005). In amphibians, either males or females care for the offspring (Reynolds et al., 2002; Furness & Capellini, 2019; Vági et al., 2019), whereas in reptiles, female-only care is widespread, but biparental care also occurs (Reynolds et al., 2002; Balshine, 2012; Halliwell et al., 2017).

Sex ratios have been suggested to play a crucial role in explaining the diversity of parental care patterns (Emlen & Oring, 1977; Kokko & Jennions, 2008; 2012; Liker et al., 2013; Székely et al., 2014). Traditionally, the operational sex ratio (OSR, hereafter defined as the proportion of males among those individuals in the population that are ready to mate) was considered a prime determinant of parental roles (Emlen & Oring, 1977). The underlying idea is that there is a trade-off between parental efficiency and competitive ability on the mating market (Magrath & Komdeur, 2003). As the members of the majority sex on the mating market have to compete more intensely for matings, these members should, so the argument goes, invest relatively more in their competitiveness than in parental care (Clutton-Brock & Parker, 1992; Kvarnemo & Ahnesjo, 1996; Simmons & Kvarnemo, 2006; Janicke & Morrow, 2018). This, in turn, induces the members of the limiting sex to be choosy and to provide more parental care. However, this argument has been criticized for three reasons. First, the existence of a universal trade-off between parental efficiency and competitive ability on the mating market is debatable (Stiver & Alonzo, 2009). In some organisms, it is likely that armament and ornamentation increasing success on the mating market are a handicap when it comes to parental care (e.g., Duckworth et al., 2003; Mitchell et al., 2007). Conspicuous colouration, for example, may attract predators to the nest (Huhta et al., 2003; Morehouse & Rutowski, 2010). However, many structures and signals that are relevant for mating do not necessarily interfere with parenting (e.g., because they are only expressed during the mating period). Second, selection on competitiveness does not necessarily lead to a high investment in competitive structures in all members of the competing sex (Baldauf et al., 2014). Mating competition may become prohibitively costly, to the point where it becomes advantageous to focus on other reproductive activities (e.g., parental care) (Kokko & Jennions, 2008; Baldauf et al., 2014). This can result in a self-reinforcing process: individuals that are less competitive on the mating market cannot expect many future matings; accordingly, they should invest a lot into each of the few matings (and the resulting brood) that they can realise. In other words, one would expect considerable variation in the level of parental care in the majority sex, and it is not self-evident that this sex should care less on average. Third, the parental care pattern has immediate repercussions on the OSR (Székely et al., 2000; Kokko & Jennions, 2008; Jennions & Fromhage, 2017). If one sex does all the caring, the availability of this sex on the mating market will typically be reduced. In other words, the causal relationship between OSR and the pattern of parental care is reciprocal (Székely et al., 2000; Kokko & Jennions, 2008): the OSR may be a “driver” of parental sex roles, but at the same time it is also “driven” by the parental care pattern. Therefore, the role of OSR as a driver of parental sex roles is, at best, ambiguous.

Recently, the adult sex ratio (ASR, defined as the proportion of males among the adult individuals in the population) has garnered considerable attention from both empiricists and theorists (Kokko & Jennions, 2008; Liker et al., 2013; Székely et al., 2014; Fromhage & Jennions, 2016; Schacht et al., 2017). Their work suggests that a male-biased ASR promotes males to provide more parental care, while a female-biased ASR leads to female-biased care. For two reasons ASR variation is considered a better predictor of parental patterns than OSR variation. First, the “Fisher condition” is applicable to the ASR, rather than the OSR. According to Fisher (1930), in diploid sexually reproducing organisms each offspring has one father and one mother. As a result, the total number of offspring produced by each sex must be equal. Any bias in the ASR has therefore a straightforward implication: the minority sex, on average, produces more offspring than the majority sex (Queller, 1997; Houston & McNamara, 2002). Thus, the more abundant sex receives less fitness revenue from mating than the rarer sex and, all other things being equal, the members of the majority sex benefit more from devoting time and energy to parental care. Second, the ASR is determined by a variety of factors (e.g., sex differences in maturation, survival, dispersal, and migration (Székely et al., 2014; Ancona et al., 2020) that are often only loosely related to parental behaviour. Accordingly, the causality between the sex ratio and the parental care pattern is expected to be reciprocal to a smaller extent in case of the ASR than in case of the OSR. However, it is important to realise that also the ASR is affected by reproduction-related feedbacks. Such feedbacks easily arise when mortality rates differ between the mating and the caring stage and/or between the sexes (Fromhage & Jennions, 2016; Jennions & Fromhage, 2017). Accordingly, it may be difficult for both, the ASR and the OSR, to disentangle cause and effect when discussing the relationship between parental care and the sex ratio.

In view of these intricacies, it is difficult, if not impossible, to predict evolutionary outcomes by using only verbal arguments. Therefore, mathematical models have been built in an attempt to understand in what way sex ratios influence parental care patterns. To be mathematically tractable, early models (e.g., Clutton-Brock & Parker, 1992; Yamamura & Tsuji, 1993; Queller, 1997; Houston & McNamara, 2002) incorporated sex ratios as a fixed parameter; accordingly, they could not address the feedbacks mentioned above. This changed with the landmark paper of Hanna Kokko and Mike Jennions (2008), who developed a simple and elegant framework to address the joint evolution of parental roles and sex ratios. Based on this framework, Kokko and Jennions (2008) drew some interesting conclusions, but as pointed out later by Lutz Fromhage and Mike Jennions (2016), their analysis is flawed due to a mistake in their fitness function. For example, Kokko and Jennions (2008) had argued that the sex that is overrepresented in the OSR should provide more care, and egalitarian biparental care should evolve in the limiting case of no differences between the sexes. In contrast, Fromhage and Jennions (2016) concluded that an OSR bias does not select for a care bias; in the limiting case of no sex differences their fitness gradient method does not predict the evolution of egalitarian care, but rather evolution to a neutral line of equilibria, ranging from male-only care via egalitarian care to female-only care. Kokko and Jennions (2008) also made the prediction that the ASR has a direct role in driving parental sex roles: according to their analysis, the more common sex in the adult population is selected to provide more care. Based on their improved fitness function, Fromhage and Jennions (2016; see also Jennions & Fromhage, 2017) showed that this conclusion is incorrect, because the route by which the ASR becomes biased may play a crucial role for the outcome of parental care evolution. In other words, parental sex roles are not driven by an ASR bias, but by the factors (e.g., sex-differential mortalities) underlying this bias. According to Jennions and Fromhage (2017), one of these factors is the maturation sex ratio (MSR, defined as the proportion of males among those individuals that are at the start of their adult life). They argue that the more common sex at maturation is selected to provide more care, and that, accordingly, MSR has the property ascribed by Kokko and Jennions (2008) to the ASR.

In the present study, we complement the mathematical analysis of Fromhage and Jennions (2016) by individual-based evolutionary simulations that make use of a very similar model structure. Such a simulation approach is important for at least three reasons. First, the fitness considerations underlying the mathematical analysis of sex role evolution are intricate and therefore error-prone. This is illustrated by the fact that the analysis of some foundational studies on the evolution of parental care is fundamentally flawed (see Houston & McNamara, 2005; Fromhage & Jennions, 2016). It is therefore useful to check the analytical predictions by means of an independent approach, which is, as our simulations, not based on the analysis of a fitness function. Second, mathematical analyses are restricted to highly simplified scenarios, as the limitations of analytical tractability are soon reached in models of sex role evolution. These limitations do not apply to simulation models. For example, sexual selection can be incorporated in a more natural way than in the framework of Kokko and Jennions (2008). Third, and most importantly, the mathematical analysis is often based on (hidden) assumptions that are not always justified. For example, the selection gradient approach used by Kokko and Jennions (2008) and Fromhage and Jennions (2016) implicitly assumes that the male and female parts of the population are monomorphic. As shown by Long and Weissing (2020), this assumption is not justified: when the sexes have conflicting interests (as in sex role evolution), the population undergoes periods of divergent selection, leading to polymorphism. Even if polymorphism is transient, it can be decisive for the evolutionary outcome. This is illustrated by the baseline model of Fromhage and Jennions (2016) (no sex differences in mortality, no parental synergism): while the selection gradient approach predicts a selectively neutral line of equilibria, the simulations reveal that there are actually two stable outcomes, either male- or female-biased care. Simulations are therefore an important check of whether the analytical methods predict the evolutionary outcome correctly.

Here, we use the simulation model of Long and Weissing (2020) to systematically study the joint evolution of parental behaviour and sex ratios (MSR, ASR, OSR) for various types of sex difference in life history parameters (maturation rate, juvenile mortality, mortality in the mating phase, mortality in the caring phase). First, we consider the case that the parents have an additive effect on offspring survival. As the mathematical model is degenerate in this case (exhibiting a neutral line of equilibria), no analytical predictions are available for this case. Second, we extend the analysis to parental synergy. By rescaling the model of Fromhage and Jennions (2016), we can systematically compare the simulation outcomes with their analytical predictions.

Throughout, we address the following questions: Do sex differences in life history characteristics have a predictable outcome on the evolution of parental care biases? How do sex ratios co-evolve with parental care patterns? Is there a consistent relationship between the bias in one of the sex ratios (MSR, ASR, OSR) and the parental care bias? In addition, we will touch upon questions as: Is, for a given parameter combination, the evolutionary outcome unique or are there alternative stable states? Does parental synergy lead to strongly different outcomes than scenarios where parental effects on offspring survival are additive? To what extent do the simulation outcomes confirm the analytical predictions of analogous mathematical models.

## Methods

The individual-based simulations were based on (a slightly simplified version of) the model of Long and Weissing (2020). We consider a randomly mating population with overlapping generations. The time structure of the model is discrete; a time unit is thought to represent one day. The model considers the evolution of two strategic parameters: *T*_*f*_ and *T*_*m*_, the number of days invested in the care of the current brood when being the female or male parent, respectively. *T*_*f*_ and *T*_*m*_ are natural numbers that are encoded on two unlinked gene loci that are expressed in a sex-specific manner. They evolve via mutation and selection, and the evolutionary outcome determines the parental sex roles in the population. For simplicity, the individuals in our model are haploid.

### Life cycle

Each day, the individuals in our model are in one of three states (see Fig. 1): the juvenile state, in which newly born offspring stay until maturation; the mating state, in which males and females seek for mating partners; and the caring state, in which individuals provide parental care to their offspring. Offspring surviving the parental care period enter the juvenile state, where they experience the sex-specific mortality rate (i.e., the probability to die per day) 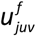 or 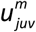, respectively. Surviving juveniles mature after *J* ^*f*^ or *J*^*m*^ days; after this time, they begin their adult life in the mating state. While in the mating state, the individuals are exposed to the sex-specific mortality rates 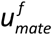 and 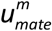. Each individual in the mating state has one mating opportunity per day: pairs are formed at random, until only one sex is left in the mating state. Unmated individuals stay in the mating state for another day. The mated individuals switch to the caring state, where they stay for *T*_*f*_ or *T*_*m*_ days, depending on their inherited parental-care strategy. The mortality rates in the caring state are 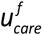 and 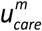, respectively. After leaving the caring state, surviving parents enter the mating state again. Each mated pair gives birth to one offspring, whose survival depends on the amount of care provided by the two parents (see below). Surviving offspring enter the juvenile state.

**Figure 1.**
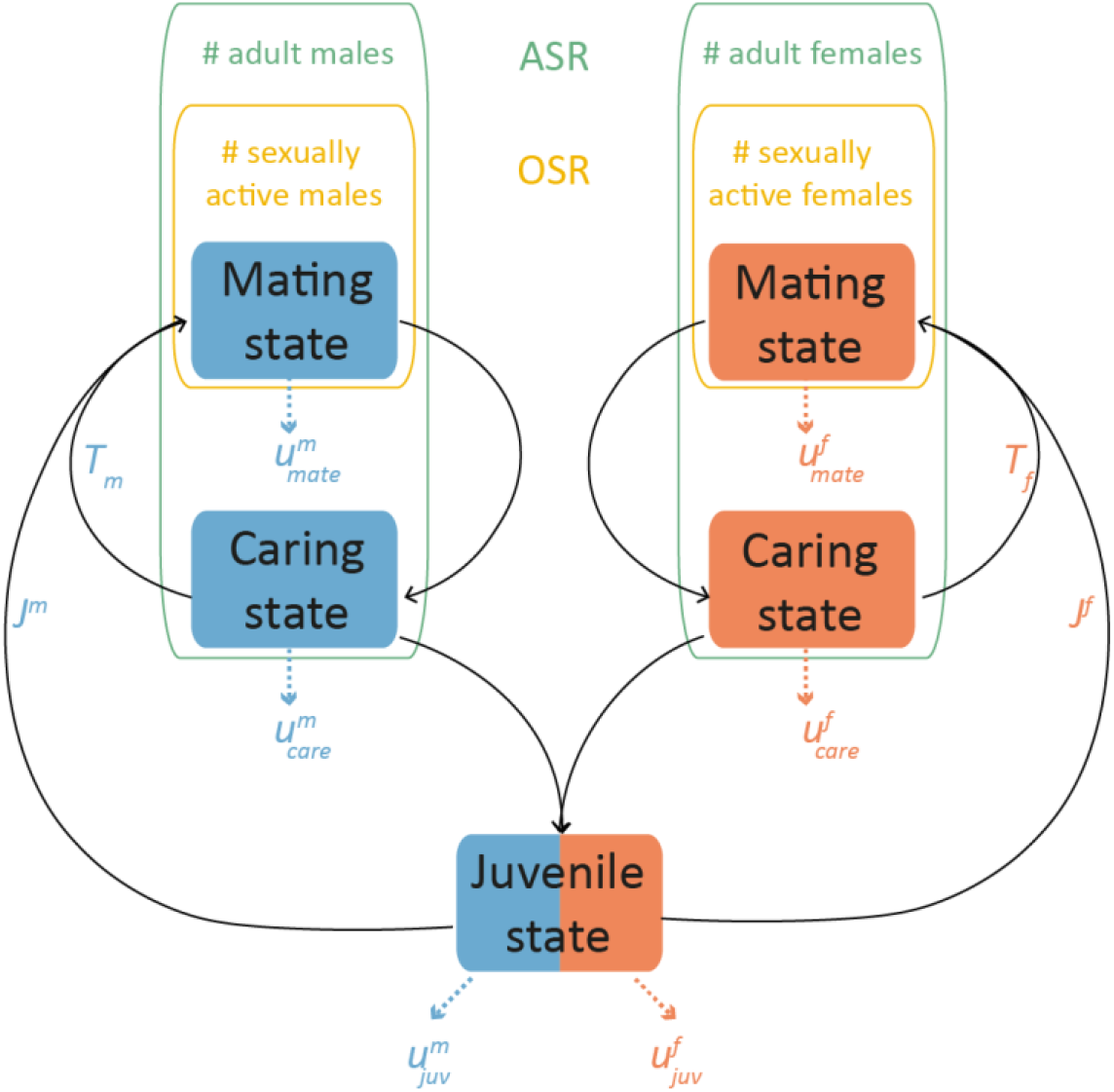
A schematic representation of the life cycle in our model. Individual males (blue) and females (red) can be in three states. Surviving offspring first spend some time in the juvenile state. After a sex-specific maturation time (*J*^*m*^ or *J*^*f*^) they enter adulthood, where they switch between the mating state and the caring state. Unless stated otherwise, the maturation time is 20 time unites for both sexes (*J*^*m*^ = *J* ^*f*^ = 20). The time in the mating state depends on the availability of mates and hence on the operational sex ratio (OSR), the proportion of males among the individuals in the mating state. In contrast, the adult sex ratio (ASR) refers to the proportion of males among all adult individuals. After mating, the mates spend *T*_*m*_ and *T*_*f*_ time units in the caring state; afterwards they return to the mating state. *T*_*m*_ and *T*_*f*_ are heritable traits that evolve subject to mutation and selection. A longer care time increases the survival probability of the offspring. In all states, (sex-specific) mortality occurs with probability 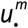 and 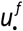 per time unit (juvenile state: 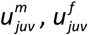, mating state: 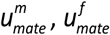, caring state: 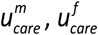). Unless stated otherwise, the mortality rates are 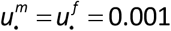, corresponding to a life expectancy of 1,000 time units.

As our main focus, we investigate the effects of sex differences in the mortality rates in each of the three states and in maturation time. In all cases, we take the following baseline scenario as our point of departure: the maturation time of both sexes is *J* ^*f*^ = *J*^*m*^ = 20 days, and the mortality rate is 0.001 day^-1^ in each state for both sexes. This means that the life expectancy of an individual in the baseline scenario is 1000 days, which, for simplicity, we regard as a proxy for generation time. In the sex-specific life-history scenarios, males and females may die at different rates or may have different maturation times. We consider one life-history parameter at a time. When we consider mortality differences in one of the states, the sex with the lowest mortality dies at the default rate of 0.001 day^-1^, while the mortality rate of the vulnerable sex ranges from 0.001 day^-1^ to 0.1 day^-1^. The mortality rate in the two other rates has the default value 0.001 day^-1^ for both sexes. In the case where juvenile females and males mature at different rates, one of the two sexes only requires 5 days to mature, while the maturation time of the other sex ranges from 5 to 50 days (with the exception of Figure S2(b)). In this scenario, all mortality rates are fixed at the default value 0.001 day^-1^.

In our model, the sex ratios at any given day can easily be calculated by counting the number of males and females in the mating state (for the OSR), the number of adult males and females (for the ASR), and the number of juveniles that are maturating on that day (for the MSR). Sex ratios are expressed as the proportion of males among all individuals in the corresponding category.

### Reproduction

Whenever a mating pair is formed, it produces a single offspring. The sex of the offspring is assigned at random, both sexes having the same probability. Offspring survival strongly depends on the total care effort provided by the parents. This is given by *T*_*tot*_ = *T*_*f*_ + *T*_*m*_ +*σ T*_*f*_ *T*_*m*_, where *T*_*f*_ and *T*_*m*_ are the inherited care strategies of the two parents, while the term *σ T*_*f*_ *T*_*m*_ (which for a given sum *T*_*f*_ + *T*_*m*_ is largest when *T*_*f*_ = *T*_*m*_) quantifies the synergistic benefits of egalitarian care. We first consider the case *σ* = 0 (no synergy), where the parents have an additive effect on offspring survival. In addition, we consider the case *σ* = 0.2, where egalitarian care provides a relatively strong benefit. As shown by Long and Weissing (2020), egalitarian biparental care evolves if *σ* = 0.2 and the life history parameters are the same for both sexes. The care-dependent survival probability of the offspring is given by 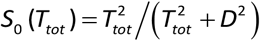, where *D* can be interpreted as the care demand of the offspring (see Long & Weissing, 2020). We chose *D* = 20 in all simulations. In addition, offspring survival is density dependent: 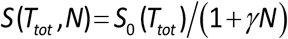, where *N* is the current total population size and *γ* is a scaling parameter. This ensures that the population size remains constant. In all simulations, we set *γ* = 0.003 ; as a result, the total population size stabilised at approximately 4000 individuals.

The offspring inherit the traits *T*_*f*_ and *T*_*m*_ from their parents. For each of the two loci, either the paternal or the maternal allele was transmitted, with equal probability. Immediately after inheritance, a mutation could occur, with probability *μ* = 0.005 per locus. In case of a mutation, the inherited allele value was either increased or decreased by 1, with equal probability. Mutations from zero to a negative allele value were neglected.

### Initialisation and replication

All simulations were started with a monomorphic population of 1000 adult males and 1000 adult females. The *T*_*f*_ -locus and *T*_*m*_ -locus were initialised at different values (see Fig. 2 and 3), The mortality rates and maturation times were varied across a wide range of parameter values, as stated above. For most parameter combinations, equilibrium was reached within 1,000 generations. In these cases, we ran 100 replicate simulations for 5,000 (or, in some cases, 50,000) generations to ensure that the results were representative. In some scenarios, there were two alternative stable outcomes. In these cases, attaining equilibrium may take a substantially longer period; therefore, we conducted 20 replicate simulations for 500,000 generations. All simulations were executed in C++.

**Figure 2.**
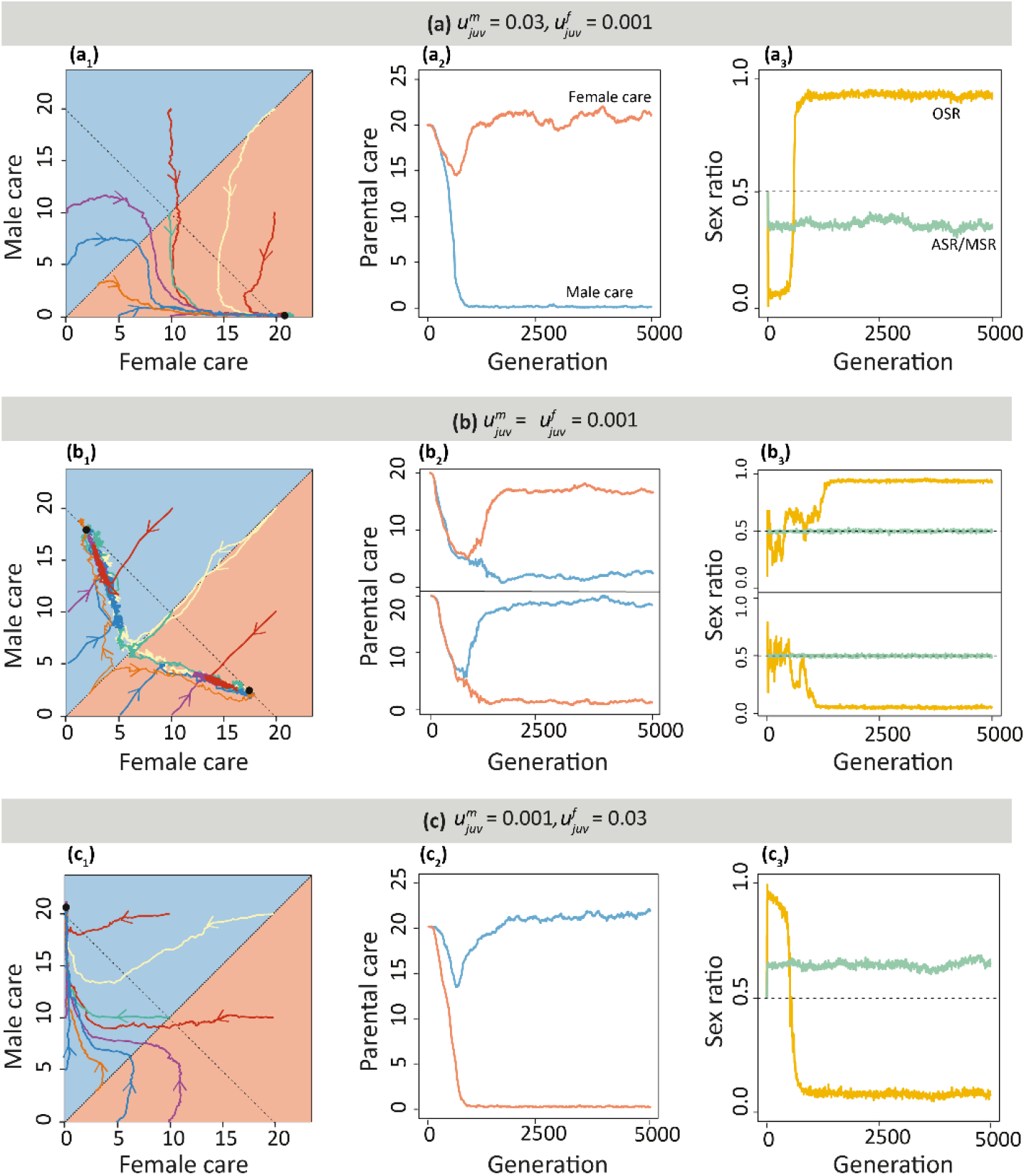
Sex differences in juvenile mortality drive sex role divergence. The graphs consider three scenarios where males and females differ only in their juvenile mortality rates (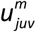 and 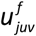). Since all adult mortality rates are the same, the ASR is identical to the MSR. **(a)** If male juvenile mortality is higher than female juvenile mortality (here: 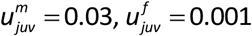), female-only care evolves, as is illustrated in (a_1_) by the coloured trajectories that, starting from different initial conditions, all converge to the point (*T*_*m*_, *T*_*f*_) = (0,20) (indicated by a black dot). The yellow trajectory, starting at high-level egalitarian care ((*T*_*m*_, *T*_*f*_) = (20,20)), is shown as a time plot in (a_2_). Panel (a_3_) shows that the ASR (=MSR) (green line) stays approximately constant at 0.35. The OSR (yellow line) is first strongly female-biased and later, when evolutionary equilibrium is attained, strongly male-biased. **(b)** In the absence of sex differences 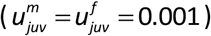, either female-biased care (upper panels in b_2_ and b_3_) or male-biased care (lower panels in b_2_ and b_3_) evolves, with equal probability. In this case, the ASR (=MSR) is unbiased and the OSR is biased toward the less-caring sex. **(c)** If female juvenile mortality is higher than male juvenile mortality (here: 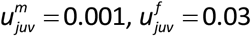), male-only care evolves, with an ASR (=MSR) and an OSR pattern that is the mirror image of the pattern in (a). In all panels, male and female mortality rates in the mating and caring state were equal to 0.001. In all simulations parents had an additive effect on offspring survival (no synergy, *σ* = 0).

**Figure 3.**
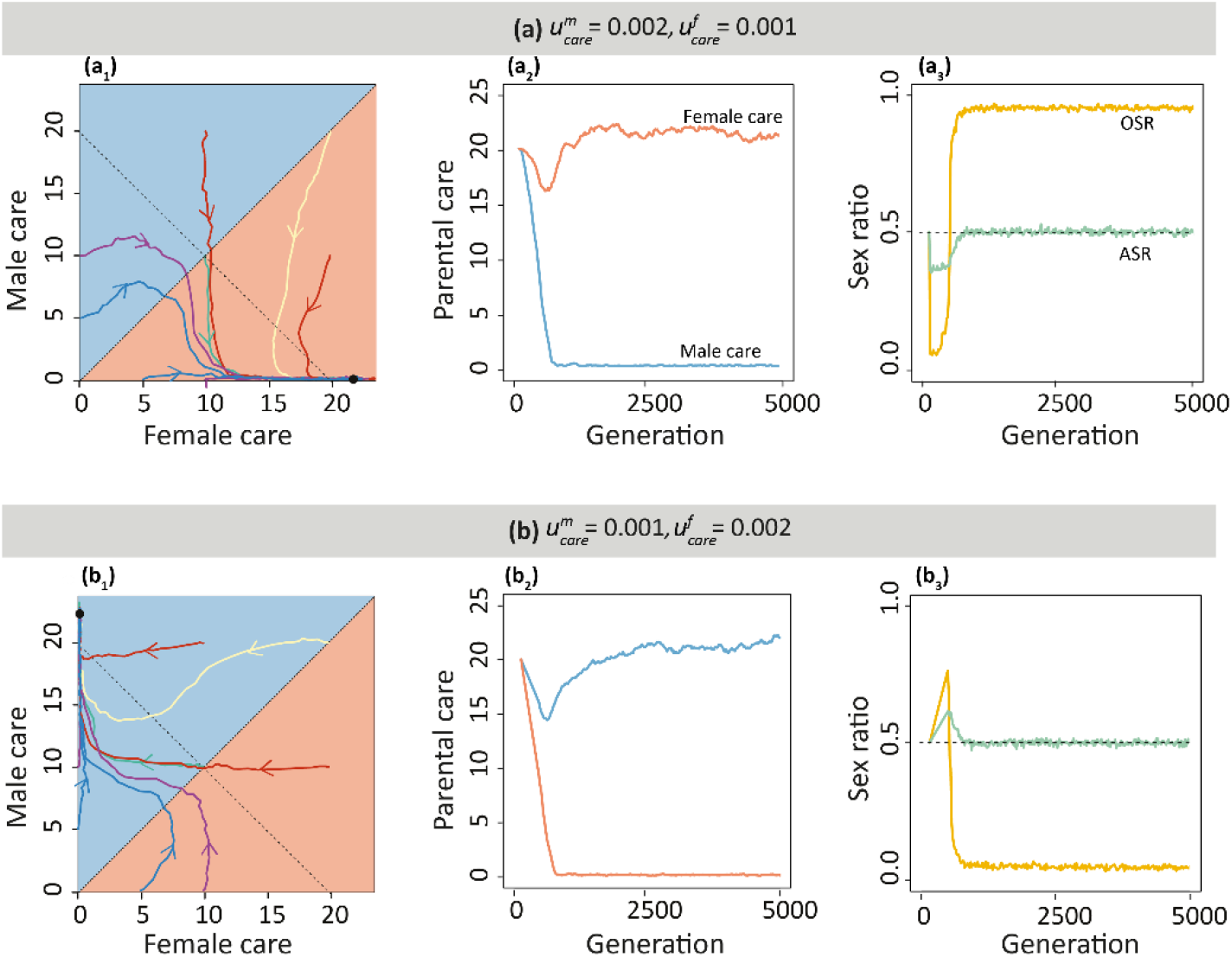
Sex differences in mortality while caring drive sex role divergence. The graphs consider two scenarios where males and females differ only in their caring mortality rates (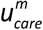 and 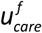). As juvenile life-history parameters are identical in this scenario, the MSR is unbiased. **(a)** If caring for offspring is more dangerous for male than for female parents (here: 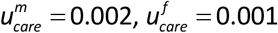), female-only care evolves, resulting in an unbiased ASR and a strongly male-biased OSR. **(b)** Male-only care evolves if females die at a higher rate during the caring stage 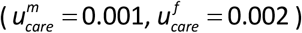, leading to an unbiased ASR and strongly female-biased OSR. Graphical conventions as in Fig. 2. Male and female mortality rates in the juvenile and mating state were all equal to 0.001. In all simulations parents had an additive effect on offspring survival (no synergy, *σ* = 0).

## Results

We first discuss the joint evolution of parental care patterns and sex ratios for the case that the parents have an additive effect on offspring survival (*σ* = 0). Later we discuss the effects of parental synergy.

### Sex differences in juvenile mortality or juvenile maturation time

Figure 2 illustrates how sex differences in juvenile mortality affect the evolution of sex-specific parental care patterns and the associated evolution of the OSR and the ASR (in this scenario, where adult mortalities are the same in both sexes, the ASR is identical to the MSR). When juvenile mortality differs between the sexes (and all other life-history parameters are the same for both sexes), uniparental care evolves, irrespective of the initial conditions (Fig. 2(a_1_,c_1_)). As illustrated in the time plots in Fig. 2(a_2_,c_2_), in evolutionary equilibrium all the care is provided by the sex with the lower juvenile mortality. In these example simulations, which started at a high degree of egalitarian biparental care (*T*_*f*_ = *T*_*m*_ = 20), the care level in both sexes first declines, as long as the total care provided (*T*_*f*_ + *T*_*m*_) exceeds the care demand of the offspring (*D* = 20). The decline in care level continues in the sex with higher juvenile mortality, until the members of this sex do not care anymore. When the total care level *T*_*f*_ + *T*_*m*_ has dropped below the care demand *D*, the sex with lower juvenile mortality increases its parental effort again, until this sex satisfies the care demands of the offspring on its own. Not surprisingly, the ASR is biased toward the sex with lower juvenile mortality (Fig. 2(a_3_,c_3_)) throughout the whole evolutionary trajectory. In contrast, the OSR changes quite dramatically in the course of evolution. In the initial period (the first 500 generations), when both sexes provide similar levels of care, the OSR is strongly skewed in the same direction as the ASR (toward the sex with lower juvenile mortality), but once uniparental care evolves, it becomes extremely skewed in the opposite direction, toward the non-caring sex. Fig. 2(b) shows the border case where *all* mortality rates are the same for both sexes. In line with our earlier study (Long & Weissing, 2020), all simulations converged to one of two evolutionary equilibria, corresponding to either strongly female-biased care and strongly male-biased care (Fig. 2(b_1_,b_2_)). The ASR (= MSR) remains unbiased, and the OSR is strongly biased toward the non-caring sex (Fig. 2(b_3_)). Figure 2 shows the evolutionary outcome for three particular combinations of mortality parameters. We will show later (see Fig. 5(a)) that these simulations are representative for a more general pattern.

In the Supplement (Figs. S1 and S2(a)), we show how the evolutionary outcome is affected by a sex difference in juvenile maturation times. Again, uniparental care or biparental care with a strong care bias evolved in all simulations. Typically, the faster maturating sex is the one that provides all or most of the care at evolutionary equilibrium (Fig. S1). However, even in case of a strong asymmetry in maturation rates there are two stable equilibria, and the opposite pattern of parental sex roles (where the more slowly maturating sex does most of the caring) evolves in a considerable percentage of the simulations (see Fig. S2(a_1_)). The ASR (which is equal to the MSR in this scenario) remains even, unless the asymmetry in maturation rates is very strong (Fig. S2(a_2_)). The OSR is, as before, strongly biased to the non-caring sex (Fig. S2(a_3_)).

We can conclude that sex differences in the juvenile state have a predictable effect on the evolution of parental care: they lead to pronounced parental sex roles, where the sex that matures faster or is exposed to lower mortality is typically, but not always, does (most of) the caring. Notice, however, that the ASR (= MSR) tends to be unbiased in case of sex differences in maturation, while it is biased in case of sex differences in mortality. Accordingly, ASR and MSR are, on their own, not sufficient to predict the outcome of parental sex role evolution.

### Sex differences in caring mortality

When the sexes differ in their mortality rates during parental-care periods, pronounced parental sex roles evolve in all simulations (Fig. 3). Irrespective of the initial conditions, the sex with the lower caring mortality tends to do the caring (Fig. 3(a_1_,b_1_)). The evolutionary trajectories of sex-specific care levels (Fig. 3(a_2_,b_2_)) show a very similar pattern as in Fig. 2. The same holds for the OSR (Fig. 3(a_3_,b_3_)), which again is strongly biased to the sex with lower mortality in the initial period (first 500 generations), where both parents are caring, and later switches to a strong bias toward the higher-mortality sex that, once evolutionary equilibrium is reached, refrains from caring. In contrast to Fig. 2, the ASR is biased toward the sex with lower caring mortality in the initial period; at evolutionary equilibrium it becomes unbiased again, because the vulnerable sex avoids the risky task of caring for the offspring (Fig. 3(a_3_,b_3_)). Figure 3 shows the evolutionary outcome for two particular mortality scenarios. We will show later (see Fig. 5(b)) that these simulations are representative for a more general pattern. However, when the sex difference in caring mortality is small, an appreciable percentage of the simulations converged to the opposite equilibrium, where parental care is provided by the sex with the *higher* caring mortality. In these cases, both ASR and OSR are biased toward the low-mortality and low-caring sex throughout the whole evolutionary trajectory (see Fig. 5(b)).

### Sex differences in mating mortality

Pronounced parental sex roles also evolve when the sexes differ in their mortality rates during the mating state (Fig. 4). Again, the sex with the lower mortality from mating typically most or all of the caring at evolutionary equilibrium (top panels in Fig. 4(a_1_,b_1_)). In this case, the ASR is biased to the caring sex (which has a lower mortality), while the OSR is biased toward the non-caring sex (top panels in Fig. 4(a_2_,b_2_)). However, even in case of considerable differences in mating mortality, two alternative evolutionary outcomes exist and a considerable percentage of all simulations end up in parental sex roles where most of the caring is done by the sex with the higher mortality during the mating state (bottom panels in Fig. 4(a,b)). When this happens, the ASR tends to be unbiased, while the OSR is strongly biased toward the non-caring sex (bottom panels in Fig 4(a_2_,b_2_)). As shown in Fig. 5(c), the outcome of the simulations in Fig. 4 is quite representative. The middle panel in Fig. 5(c) seems to indicate a considerable bias in the ASR in those cases where the high-mortality sex does the caring. However, this is most likely reflects the fact that non-equilibrium periods are included in Fig. 5(c), due to the lower stability of this sex role pattern (as indicated by the bottom panels of Fig. 4(a,b)).

**Figure 4.**
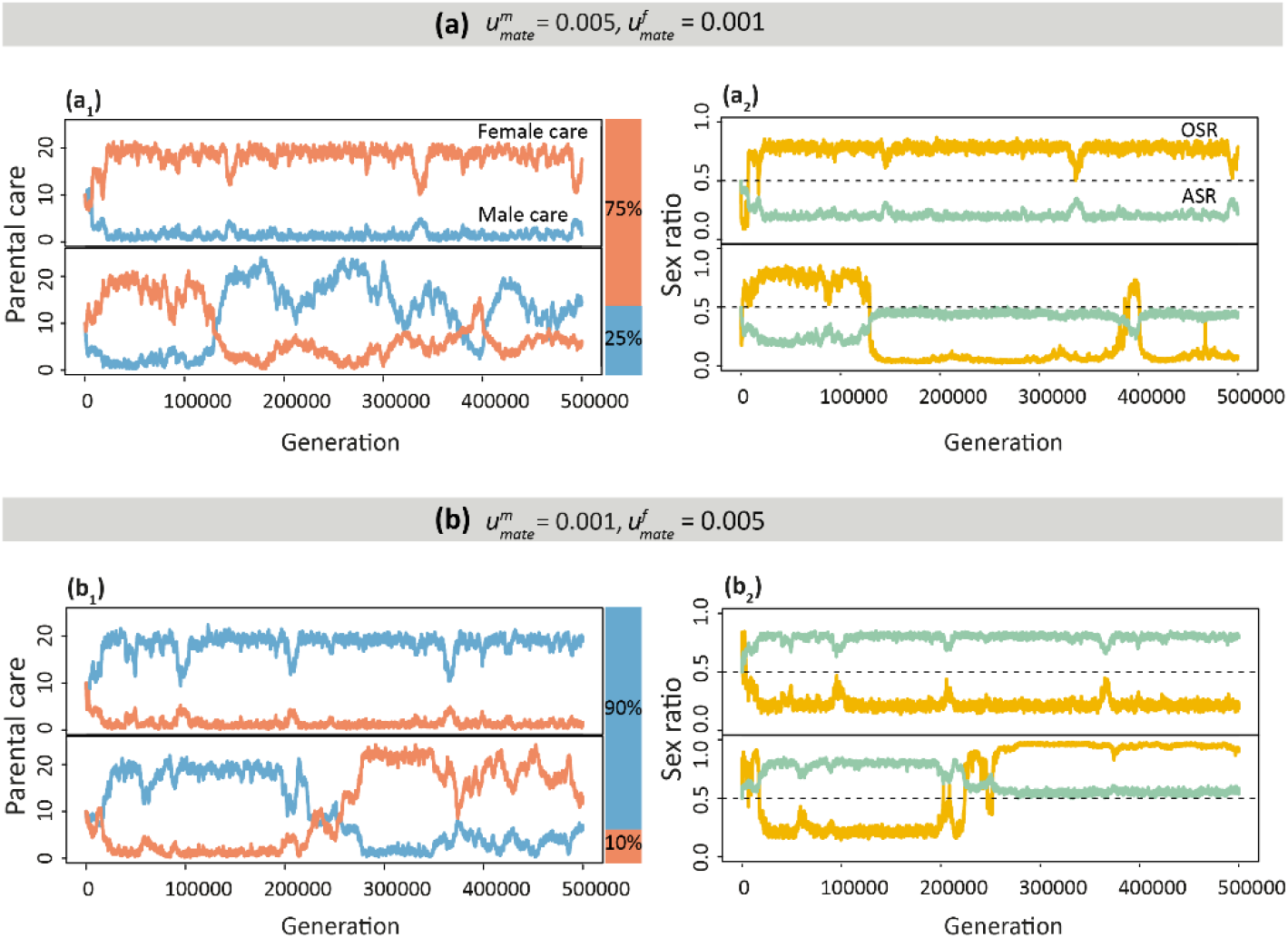
Sex differences in mortality while competing for mates drive the evolution of parental roles. The graphs consider two scenarios where males and females differ only in mortality when they are in the mating state (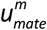 and 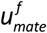). As juvenile life-history parameters are identical in this scenario, the MSR is unbiased. **(a)** When males face greater risk when competing for mates 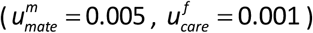, female-biased care evolved in 15 out of 20 replicate simulations (75%; upper panels in (a_1_) and (a_2_)), corresponding to a strongly female-biased ASR and a strongly male-biased OSR. In the other 25% of the simulations there are extended periods of male-biased care (lower panels); in these periods, the ASR is slightly female-biased, while the OSR is strongly female-biased. **(b)** When females face a higher mortality risk when in the mating state 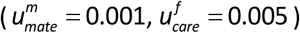, male-biased care evolved in 18 out of 20 replicate simulations (90%; upper panels in (b_1_) and (b_2_)), associated with a male-biased ASR and a female-biased OSR. In the other 10% of the simulations there are extended periods of female-biased care, associated with an ASR that is slightly male-biased and a strongly male-biased OSR. Male and female mortality rates in the juvenile and caring state were all equal to 0.001. 20 replicate simulations were run for 500,000 generations per parameter setting. In all simulations parents had an additive effect on offspring survival (no synergy, *σ* = 0).

**Figure 5.**
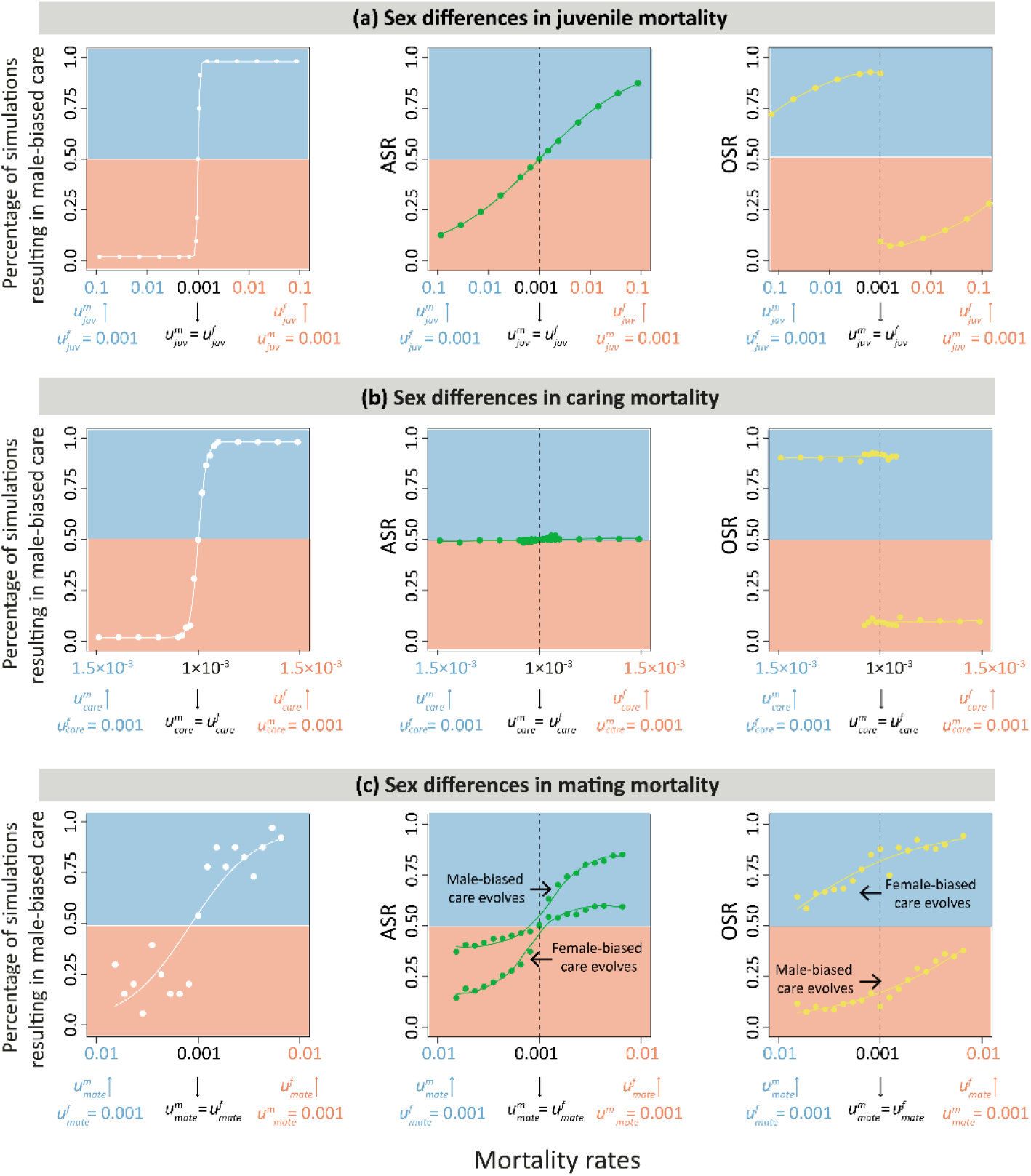
Implications of sex differences in mortality on the joint evolution of parental roles and sex ratios in the absence of parental synergy. Outcome of a large number of simulations considering sex differences in **(a)** juvenile mortality (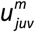 and 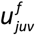), **(b)** mortality while caring for the offspring (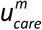and 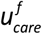), and **(c)** mortality while competing for mates (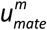 and 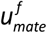). In (a) and (b), each dot represents 100 replicate simulations run for 5,000 generations for the mortality parameters indicated on the horizontal axis of each panel; in (c), each dot represents 20 replicate simulations run for 500,000 generations. All simulation started from egalitarian care (*T*_*m*_ = *T*_*f*_ = 10). The left panels show the percentage of simulations resulting in male-biased care. In all simulations, pronounced parental sex roles evolved, where one sex does most of the caring. In all three mortality scenarios, the sex with the lowest mortality tended to do most of the caring. The alternative outcome (that the sex with highest mortality does most of the caring) did occur in scenarios (a) and (b), but only when sex disparities in mortality rates were very small. In contrast, this alternative outcome arose more frequently in scenario (c). The middle and right panels show the ASR and OSR averaged over the simulations; in scenario (c) ASR and OSR are shown separately for the cases where male-biased care evolved and the cases where female-biased care evolved. In all cases, the OSR is biased toward the non-caring sex. In (a) and (c), the ASR is biased toward the low-mortality sex, while an even ASR evolves in scenario (b). In all simulations parents had an additive effect on offspring survival (no synergy, *σ* = 0).

### Overview of evolutionary outcomes in the absence of parental synergy

As shown in the overview figures (Fig. 5 and Fig. S2(a)) and in Table 1, a clear pattern arises in the no-synergy scenario (*σ* = 0) considered thus far, where the parents have an additive effect on offspring survival. Pronounced parental sex roles evolved in all simulations, in a predictable manner. Generally, the sex with the faster maturation or the lower mortality tends to become the caring sex, unless the sex difference in maturation times or mortalities is very small. In the latter case, two alternative stable outcomes, corresponding to either strongly male-biased care or strongly female-biased care do exist, as analysed in detail in our earlier study (Long & Weissing, 2020). However, there are two scenarios where alternative outcomes also occur in case of considerable asymmetry between the sexes: when the sexes differ in maturation time (Fig. S2(a)), and when they differ in mortality during the mating phase (Fig. 5(c)). In these two scenarios, the opposite outcome (that parental care is strongly biased toward the sex with slow maturation or with high mortality) also evolves in a considerable percentage of the simulations.

**Table 1.** Overview of the effects of sex differences in life-history characteristics on the joint evolution of parental roles and sex ratios. The table summarises the conclusions of our simulation study for two scenarios: (**a**) the parents have an additive effect on offspring survival (no synergy, *σ* = 0); (**b**) the parents have a synergistic effect on offspring survival (*σ* = 0.2). When the sexes differ in mortality in one of the life history states, the sex with a lower mortality dies at a rate of 0.001 day^-1^, while the other sex dies at a higher rate. The mortality rates in the other states were set to 0.001 day^-1^ for both sexes. When the sexes mature at different rates, the faster-maturing sex takes 5 days to mature, while the slower-maturing sex takes longer. In this care, all mortality rates were fixed at 0.001 day^-1^.

| (a) Parents have an additive effect on offspring survival |  |  |  |  |
| --- | --- | --- | --- | --- |
| Sex differences in: | Parental care pattern | MSR | ASR | OSR |
| Juvenile mortality | The low-mortality sex does most of the caring | Biased toward the high-care sex | Biased toward the high-care sex | Biased toward the low-care sex |
| Maturation rate | The fast-maturing sex typically, but not always, does most of the caring | Unbiased | Unbiased | Biased toward the low-care sex |
| Caring mortality | The low-mortality sex does all of the caring | Unbiased | Unbiased | Biased toward the non-caring sex |
| Mating mortality | The low-mortality sex typically, but not always, does most of the caring | Unbiased | Biased toward the low-mortality sex | Biased toward the low-care sex |
| (b) Parents have a synergistic effect on offspring survival |  |  |  |  |
| Sex differences in: | Parental care pattern | MSR | ASR | OSR |
| Juvenile mortality | Both parents care, with the low-mortality sex doing most of the caring | Biased toward the high-care sex | Biased toward the high-care sex | Biased toward the low-care sex |
| Maturation rate | Both parents care, with the fast-maturing sex doing most of the caring | Unbiased | Unbiased | Biased toward the low-care sex |
| Caring mortality | A small sex difference in caring mortality leads to a strongly sex-biased or even to uniparental care, with the low-mortality sex doing most/all of the caring | Unbiased | Unbiased | Biased toward the non-caring sex |
| Mating mortality | Egalitarian biparental care | Unbiased | Biased toward the low-mortality sex | Biased toward the low-mortality sex |

In many of the scenarios considered, the evolved sex roles were associated with a bias in the ASR: the sex that does the caring is overrepresented in the population. This, however, is not always the case. When parental sex roles evolve in response to sex differences in maturation time (Fig. S2(a_2_)) or in caring mortality (Fig. 5(b), middle panel), the ASR remains unbiased, despite pronounced sex differences in parental care. When the sexes differ in mating mortality (Fig. 5(c), middle panel), female-only care can also evolve in case of a male-biased ASR, while male-only care can evolve in case of a female-biased sex ratio. In other words, a bias in the adult sex ratio is not a reliable indicator for the outcome of parental sex role evolution. In all simulations considered thus far, the OSR is always biased toward the non-caring sex. Contrary to predictions in the literature (Kokko & Jennions, 2008; Fromhage & Jennions, 2016), however, the ASR and the OSR are often not biased in the same direction and not always responding in a similar way to changes in a parameter (e.g., Fig. 5(a)).

### Overview of evolutionary outcomes in the presence of parental synergy

For all the scenarios considered above, we also ran replicate simulations for the case that parents have a synergistic effect on offspring survival (*σ* = 0.2). The results are summarised in Fig. 6, Fig. S2(b), and in Table 1. In contrast to the case of additive parental effects reported above (*σ* = 0), now biparental care evolved, unless the sex differences in mortality rates or maturation times were very large. The figures therefore do not show the percentage of simulations resulting in male-only or female-only care, but the average evolved care level in males and females. For all parameter combinations considered, all 100 replicate simulations converged to the same equilibrium outcome, and equilibrium was typically reached within a few hundred generations.

**Figure 6.**
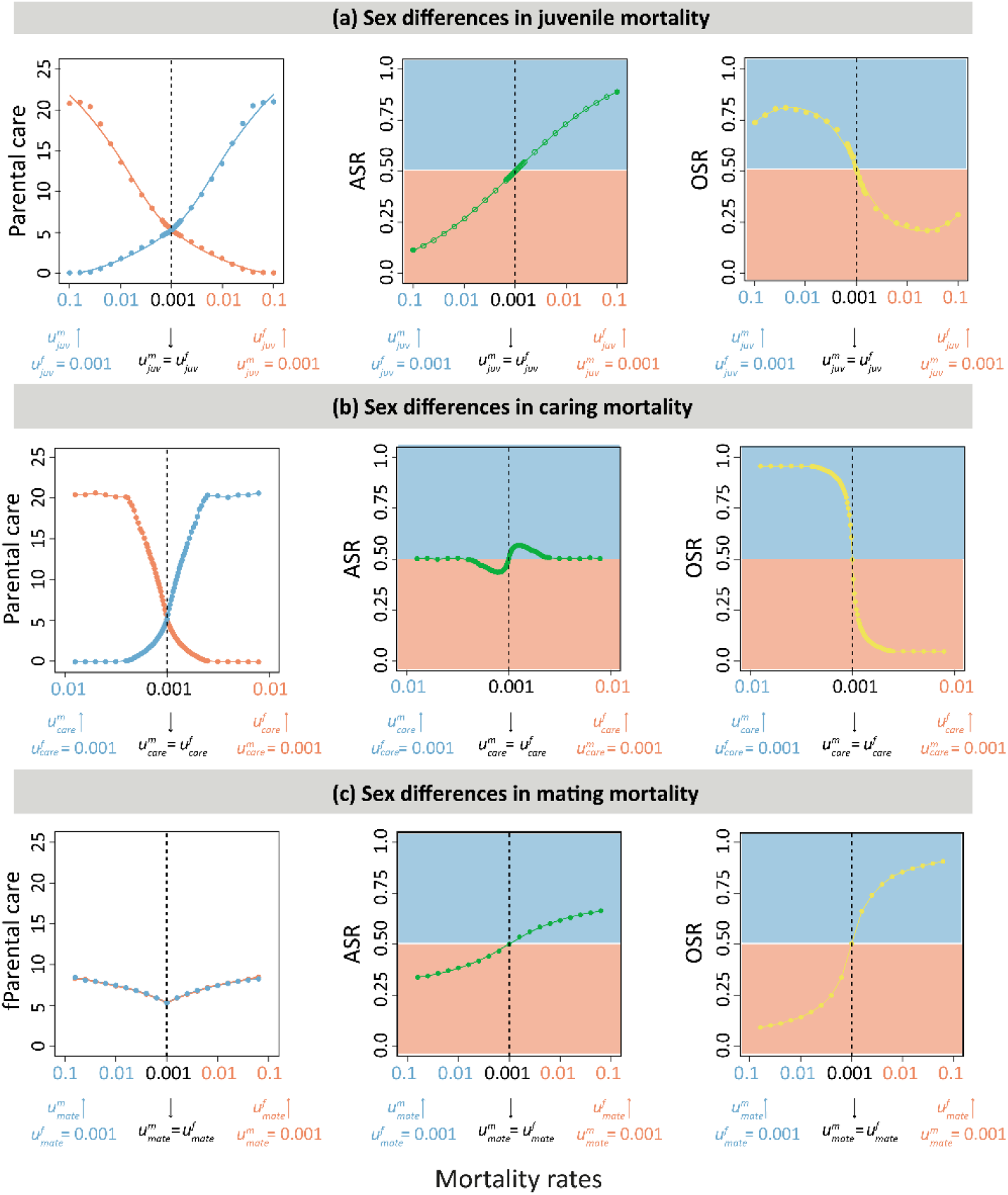
Implications of sex differences in mortality when biparental care has a synergistic effect. Outcome of a large number of simulations considering sex differences in **(a)** juvenile mortality, **(b)** mortality while caring for the offspring, and **(c)** mortality while competing for mates. In all panels, each dot represents 100 replicate simulations run for 5,000 generations (starting from egalitarian care: *T*_*m*_ = *T*_*f*_ = 10) for the mortality parameters indicated on the horizontal axis. In contrast to Fig. 5, biparental care has a synergistic effect on offspring survival (here: *σ* = 0.2). In this case, biparental care evolves when sex-specific mortalities are not too different. Therefore, the left panels now show the average level of male and female care (averaged over the 100 replicates). Notice that in scenario (c) egalitarian care evolved in all cases, even if mortality in the mating state differed strongly between the sexes.

When juvenile mortality differs between the sexes (Fig. 6(a)), both parents care, but the level of parental care is inversely related to juvenile mortality. In other words, the sex with lower juvenile mortality cares more than the sex with higher juvenile mortality. The ASR (which is identical with the MSR in this scenario) is biased toward the sex with lower juvenile mortality, which is also the sex doing most of the caring. Interestingly, the OSR shows the opposite bias than the ASR: it is biased toward the non-caring sex. Moreover, the relationship of the OSR with the degree of juvenile mortality is non-monotonic. A very similar picture arises when juvenile maturation times differ between the sexes (Fig. S2(b)): both parents care, but the level of care is positively related to maturation time. Hence, the faster maturating sex cares more than the sex with a longer maturation time. The ASR is biased toward the non-caring sex, while the OSR is biased toward the caring sex.

When the mortality rate during the period of parental care differs between the sexes (Fig. 6(b)), parental sex roles are more pronounced than in the other scenarios – already a relatively small sex difference in caring mortality leads to a strong sex bias or even to uniparental care. Again, the sex with lower caring mortality does most of the caring. For parameter combinations leading to uniparental care, the ASR is unbiased. When biparental care occurs, the ASR is biased toward the sex doing more of the caring. The opposite is the case for the OSR, which is strongly biased toward the non-caring sex.

When the mortality rate in the mating phase differs between the sexes (Fig. 6(c)), egalitarian biparental care evolves for all parameters considered. ASR and OSR are both biased toward the sex with lower mating mortality.

In case of parental synergy, our simulations can be compared with analytical predictions that make use of the selection gradient method of Fromhage and Jennions (2016). The results are shown in Fig. S3. A comparison of this figure with Fig. 6 reveals that the simulations agree very well with the predictions of the Fromhage-Jennions model (see the caption of Fig. S3 for details). This is in contrast to the model variant without parental synergy, where the two approaches led to very different conclusions (see Long & Weissing, 2020). We also run the Fromhage-Jennions model for another parametrization (Fig. S4). A comparison of Figs. S3 and S4 reveals that some of the patterns described above are not general, in that they depend on the parametrization of the model. For example, ASR and OSR show the opposite bias in Fig. S3(a), while they are biased in the same direction in Fig. S4(a) (see the caption of Fig. S4 for more details).

## Discussion

Using individual-based evolutionary simulations, we investigated the implications of sex differences in several life-history parameters on the joint evolution of parental sex roles and various sex ratios. Our conclusions are summarised in Table 1. Throughout, we considered two cases: one in which the parents have an additive effect on offspring survival and one in which they have a synergistic effect. In the first case, we consistently observed the evolution of pronounced parental sex roles (typically uniparental care), while in the second case biparental care (with a certain care bias) evolved in most simulations. Our study reveals that sex differences in life history characteristics have a systematic and predictable effect on parental care patterns. As a general rule, the sex with the lower mortality or the faster maturation is selected to provide most (or all) of the care (Figs. 5 and 6). However, in the case of additive parental effects, the opposite outcome (the sex with the highest mortality or the slowest maturation does most of the caring) evolved in a substantial proportion of the simulations (Fig. 5).

Two similar lines of reasoning, leading to opposing conclusions, can explain the different outcomes. The first line is based on biased sex ratios and explains why a lower mortality (or a faster maturation) is associated with a higher level of parental care. The argument goes like this: the sex with the lower mortality (or the faster maturation) is overrepresented in the population. In view of the Fisher condition, the members of the majority sex (here: the sex with lower mortality) have a lower *per capita* reproductive success. In our model, this means that the expected number of future matings is lower for a member of the majority sex than for a member of the minority sex. In the trade-off between current and future reproduction, the members of the majority sex should therefore place (relatively) more emphasis on the current brood than the members of the minority sex. The majority sex is therefore more strongly selected to provide care. Eventually, this asymmetry may result in parental care roles where the parent belonging to the majority sex (here: the sex with lower mortality) does most of the caring. The second line of reasoning makes use of similar life-history considerations, but arrives at the opposite conclusion. Here, the argument goes like this: the sex with the higher mortality has a shorter life expectancy and consequently a lower potential for future reproduction (Stearns, 1976; Klug et al., 2013). Using the same argument as before, this sex should be more strongly selected to invest in the current brood, leading to a parental sex bias toward the sex with higher mortality. In a situation like this, where two similar lines of reasoning lead to opposite conclusions, verbal reasoning alone is not sufficient to predict the evolutionary outcome.

What role, then, do sex ratios play in the evolution of parental care? A given sex ratio bias (be it in the OSR or the ASR) does of course affect the selection pressure on sex-specific parental care patterns. When, for example, the OSR is sex-biased (and the sexes do not differ in adult life-history traits), the members of the majority sex have a lower expected number of matings over lifetime, favouring a greater investment in parental care over mating competition (Kokko & Jennions, 2008). However, the OSR is not fixed and may change in the course of evolution (see Figs. 2 to 4). The reason is that the causality between OSR and parental sex roles is not unidirectional but reciprocal (Székely et al., 2000; Jennions & Fromhage, 2017): if one of the sexes provides most (or all) of the care, the members of that sex will typically be less available for mating, shifting the OSR toward the less-caring sex. In all scenarios explored in this study (with the exception of Fig. S4(b)), the OSR is, at evolutionary equilibrium, skewed toward the sex that provides less care. Yet, this should not lead to the conclusion that parental sex roles evolve in response to an OSR bias.

The opposite seems to be the case. As revealed by the evolutionary trajectories (e.g., Figs. 2 and 4), the OSR can change strongly in the course of evolution, switching from a strong male-bias to a strong female-bias or *vice versa*. A close inspection of the trajectories reveals that changes in the parental care patterns tends to precede the change in OSR. From this we conclude that, at least for the random-mating scenarios considered in this study, parental sex roles drive the OSR, and not the other way around.

If we consider an ASR bias in isolation, the overrepresented sex in the adult population is most strongly selected to do the caring. This is because the Fisher condition applies to the ASR (as long as it is relatively constant over individual lifetime): at any given point in time, the offspring produced at that time have one adult mother and one adult father, implying that the more common sex in the adult population has a lower *per capita* reproductive output. As argued above, this tips the balance between current and future reproduction toward a higher investment in the current offspring. Thus, individuals of the overrepresented sex in the ASR are expected to do most of the caring. In a large number of simulations, we indeed found that female-biased care evolves under a female-biased ASR, whereas male-biased care evolves under a male-biased ASR (Figs. 5(a) and 6(a)). However, we also observed that under some circumstances (e.g., sex-specific caring mortality), strongly sex-biased care can evolve in the absence of ASR bias (Figs. 5(b) and 6(b)), and that under other conditions (e.g., sex-specific mating mortality), the underrepresented sex can even be associated with a much higher level of care (Figs. 5(c) and 7). This indicates that, as in the case of the OSR, there is feedback from parental care roles to the ASR, leading to a change in the ASR over evolutionary time (see the lower panels in Fig. 5).

To illustrate the reciprocal causation between ASR and parental care, we take the most noticeable case where the sexes differ in the mortality rate while caring. In this scenario, the members of the sex with the higher mortality bear a higher ‘burden’; as a consequence, the members of the other sex are more strongly selected to care. In our model, this leads to extreme parental sex roles, where the members of the high-mortality sex abstain entirely from caring. This way, they avoid the high-mortality life-history state altogether, eventually leading to an unbiased ASR. Thus, selection on the ASR and sex roles interact in a dynamic manner, making it challenging to attribute a driving role to the ASR in the evolution of parental care. From our simulations, we arrive at the same conclusion as the analytical study of Fromhage and Jennions (2016): the processes by which the ASR becomes biased, rather than the ASR itself, causes sex role divergence.

To what extent, then, can MSR predict parental sex roles? According to the studies of Fromhage and Jennions (2016) and Jennions and Fromhage (2017), the MSR may be more relevant for the evolution of parental care than the ASR. In our simulations, we observed a clear-cut relationship between the MSR and care patterns when the MSR was biased due to sex-differential juvenile mortality: male-biased care evolved whenever the MSR was male-biased, while female-biased care evolved when the MSR was female-biased (Figs. 2 and 5(a)). This, however, is not a universal pattern, because other processes, different from juvenile mortality, also play a role. For example, when the sexes mature at different rates, uniparental male or female care evolves, despite of an unbiased MSR (Figs. S1 and S2(a)). Moreover, even strong biases in the juvenile state can be ‘overruled’ by weak biases in an adult state. This is illustrated in Fig. 7, which considers a scenario where female juveniles die at a much higher rate than male juveniles 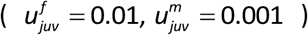, while males have a higher mortality than females while caring 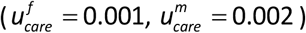. In this scenario, female-only care evolves, despite of the fact that the MSR (and the ASR, which is identical to the MSR at evolutionary equilibrium) becomes biased in favour of males. Again, we conclude that also the MSR is affected by reciprocal causality, implying that an MSR bias can change in the course of evolution (as shown in Fig. 7).

**Figure 7.**
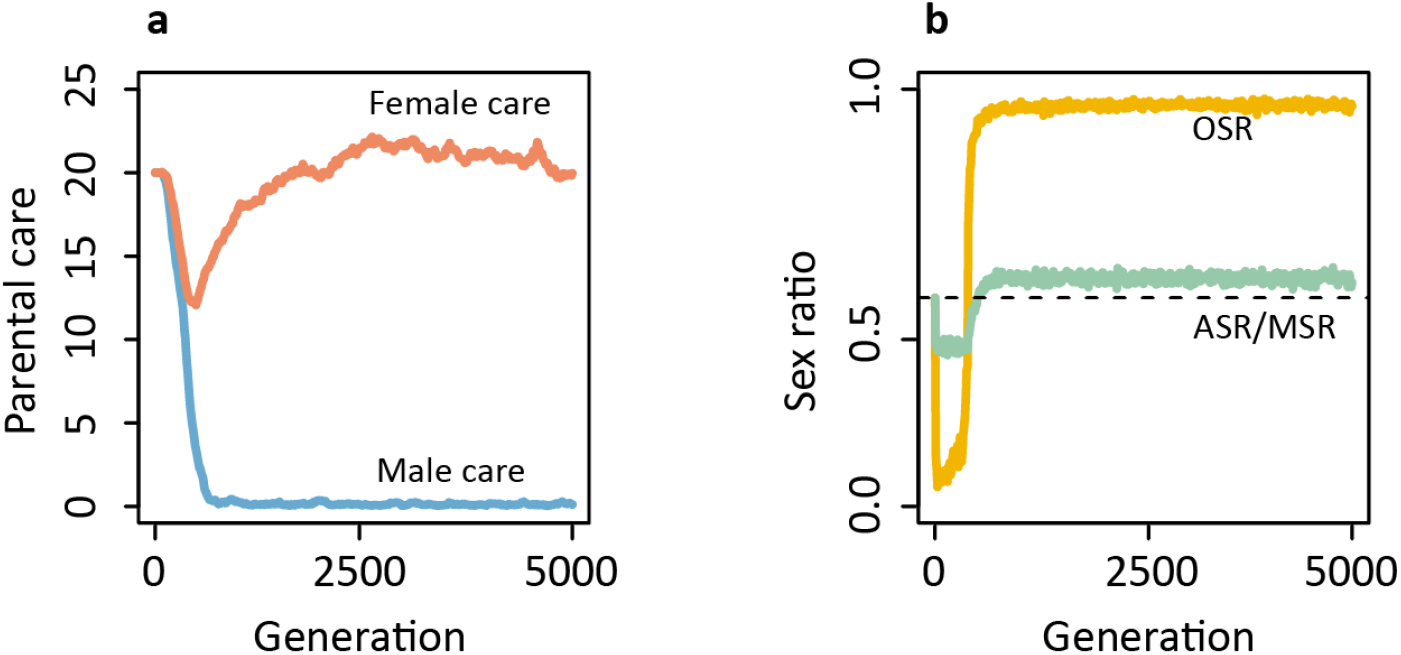
Parental sex roles are not necessarily predicted by the maturation sex ratio (MSR) or the adult sex ratio (ASR). When female juvenile mortality is higher than male juvenile mortality (here: 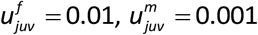), the maturation sex ratio is male-biased. In the absence of sex differences in adult mortality, this would select for male care (see Fig. 5(a)). In the simulation shown here, there is an additional mortality bias in the adult stage: males die at a higher rate than females in the caring state (here: 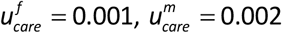). **(a)** Females are selected to care for their offspring, despite of the fact that **(b)** the MSR and the ASR are both biased in favour of males. Male and female mortality rates in the mating state were set to 0.001. Since males escape from the hazardous caring activity, adult males and adult females die at the same rate in the end, the ASR is then entirely determined by the MSR. In this simulation parents had an additive effect on offspring survival (no synergy, *σ* = 0).

Summarising all our findings, we arrive at the conclusion: none of the sex ratios should be viewed as a ‘driver’ of the evolution of parental sex roles; instead, sex disparities in life-history characteristics drive the joint evolution of sex ratios and parental sex roles. An important reason is that the sex ratios in our model are not fixed but dynamic in evolutionary time: they jointly evolve with the parental care patterns, and both the strength and the direction of the sex ratio biases often changes in the course of evolution.

Having said this, we would like to qualify our above conclusion that sex differences in life history characteristics are *the* drivers of parental sex roles. This holds in our model, where the life history characteristics are fixed by assumption. In reality, mortality rates and maturation times are also evolvable properties. For example, female preferences can induce males to develop ‘handicapping’ traits compromising their survival (Zahavi, 1975; Maynard Smith, 1991; Kuijper et al., 2012). It would therefore be interesting to consider models where the life history parameters are not externally given but at least partly ‘internalised’. In such a case, even more feedback can arise, potentially leading to different conclusions regarding the causality underlying the evolution of parental sex roles.

In our current work, we examine fairly straightforward scenarios. Most of the time, we simply look at one aspect of life history at a time. In natural systems, the sexes differ at various life stages (Orzack et al., 2015; Storchová & Hořák, 2017). The systematic investigation of the interplay of various sex differences is beyond the scope of this study. Here, we only show by means of an example (Fig. 7) that a sex difference in one mortality component (here: at the time of caring) seems to be more important than a difference in another component (here: juvenile mortality). Is this a special feature or does it reflect a general principle? What if other factors, such as sex-specific mating mortality, are introduced? Studying the interplay of such factors may seem a plausible next step, but such an endeavour may soon become unrealistic, as it would require a huge number of simulations.

For simplicity, we have focussed on random mating, thus neglecting sexual selection, which most likely will play an important role in the evolution of parental sex roles. The evolutionary interplay between sexual selection and parental care patterns may be more intricate than verbal or simple mathematical models suggest. This is exemplified in the previous study (Long & Weissing, 2020), where this interplay led to two alternative outcomes: one with strongly female-biased care associated with choosy females and bright males, and another with strongly-male biased care associated with non-choosy females and dull males. Will this interesting pattern (including long-term switches between the outcomes) remain when sex differences in life history characteristics are incorporated in the model? How will males allocate their resources to mating competition (ornamentation) on the one hand, and to parental care on the other, if ornamentation does not only affect mating success, but also mortality in the mating and the caring state? Addressing such questions seems interesting and important, but we have to leave this to a future attempt.

## Data availability

This study is theoretical; no new empirical data were generated.

## Code availability

The C++ simulation code and a Mathematica file with an implementation of the fitness gradient method are available for download from https://github.com/xiaoyanlong/sexratio.

## References

Ancona, S., Liker, A., Carmona-Isunza, M. C., & Székely, T. (2020). Sex differences in age-to-maturation relate to sexual selection and adult sex ratios in birds. Evolution Letters, 4(1), 44–53.

Baldauf, S. A., Engqvist, L., & Weissing, F. J. (2014). Diversifying evolution of competitiveness. Nature Communications, 5(1), 1–8.

Balshine, S. (2012). Patterns of parental care in vertebrates. In N. J. Royle, P. T. Smiseth & M. Kölliker (Eds.), The evolution of parental care (pp. 62–80). Oxford University Press.

Blumer, L. S. (1979). Male parental care in the bony fishes. The Quarterly Review of Biology, 54(2), 149– 161.

Clutton-Brock, T. H. (1991). The evolution of parental care. Princeton University Press.

Clutton-Brock, T. H., & Parker, G. A. (1992). Potential reproductive rates and the operation of sexual selection. The Quarterly Review of Biology, 67(4), 437–456.

Cockburn, A. (2006). Prevalence of different modes of parental care in birds. Proceedings of the Royal Society B: Biological Sciences, 273(1592), 1375–1383.

Duckworth, R. A., Badyaev, A. V., & Parlow, A. F. (2003). Elaborately ornamented males avoid costly parental care in the house finch (Carpodacus mexicanus): a proximate perspective. Behavioral Ecology and Sociobiology, 55(2), 176–183.

Emlen, S. T., & Oring, L. W. (1977). Ecology, sexual selection, and the evolution of mating systems. Science, 197(4300), 215–223.

Fisher, R.A. 1930. The genetical theory of natural selection. Oxford University Press.

Fromhage, L., & Jennions, M. D. (2016). Coevolution of parental investment and sexually selected traits drives sex-role divergence. Nature Communications, 7(1), 1–11.

Furness, A. I., & Capellini, I. (2019). The evolution of parental care diversity in amphibians. Nature Communications, 10(1), 1–12.

Halliwell, B., Uller, T., Holland, B. R., & While, G. M. (2017). Live bearing promotes the evolution of sociality in reptiles. Nature Communications, 8(1), 1–8.

Houston, A. I., & McNamara, J. M. (2002). A self–consistent approach to paternity and parental effort. Philosophical Transactions of the Royal Society of London. Series B: Biological Sciences, 357(1419), 351–362.

Huhta, E., Rytkönen, S., & Solonen, T. (2003). Plumage brightness of prey increases predation risk: an among-species comparison. Ecology, 84(7), 1793–1799.

Janicke, T., & Morrow, E. H. (2018). Operational sex ratio predicts the opportunity and direction of sexual selection across animals. Ecology Letters, 21(3), 384–391.

Jennions, M. D., & Fromhage, L. (2017). Not all sex ratios are equal: the Fisher condition, parental care and sexual selection. Philosophical Transactions of the Royal Society B: Biological Sciences, 372(1729), 20160312.

Klug, H., Bonsall, M. B., & Alonzo, S. H. (2013). Sex differences in life history drive evolutionary transitions among maternal, paternal, and bi-parental care. Ecology and Evolution, 3(4), 792–806.

Kokko, H., & Jennions, M. D. (2008). Parental investment, sexual selection and sex ratios. Journal of Evolutionary Biology, 21(4), 919–948.

Kokko, H., & Jennions, M. D. (2012). Sex differences in parental care. In N. J. Royle, P. T. Smiseth & M. Kölliker (Eds.), The evolution of parental care (pp. 102–114). Oxford University Press.

Kuijper, B., Pen, I., & Weissing, F. J. (2012). A guide to sexual selection theory. Annual Review of Ecology, Evolution, and Systematics, 43, 287–311.

Kvarnemo, C., & Ahnesjo, I. (1996). The dynamics of operational sex ratios and competition for mates. Trends in Ecology & Evolution, 11(10), 404–408.

Liker, A., Freckleton, R. P., & Székely, T. (2013). The evolution of sex roles in birds is related to adult sex ratio. Nature Communications, 4(1), 1–6.

Long, X., & Weissing, F. J. (2020). Individual variation in parental care drives divergence of sex roles. Biorxiv.

Magrath, M. J., & Komdeur, J. (2003). Is male care compromised by additional mating opportunity? Trends in Ecology & Evolution, 18(8), 424–430.

Mank, J. E., Promislow, D. E., & Avise, J. C. (2005). Phylogenetic perspectives in the evolution of parental care in ray-finned fishes. Evolution, 59(7), 1570–1578.

Maynard Smith, J. (1991). Theories of sexual selection. Trends in Ecology & Evolution, 6(5), 146–151.

Mitchell, D. P., Dunn, P. O., Whittingham, L. A., & Freeman-Gallant, C. R. (2007). Attractive males provide less parental care in two populations of the common yellowthroat. Animal Behaviour, 73(1), 165–170.

Morehouse, N. I., & Rutowski, R. L. (2010). In the eyes of the beholders: female choice and avian predation risk associated with an exaggerated male butterfly color. The American Naturalist, 176(6), 768–784.

Orzack, S. H., Stubblefield, J. W., Akmaev, V. R., Colls, P., Munné, S., Scholl, T., … & Zuckerman, J. E. (2015). The human sex ratio from conception to birth. Proceedings of the National Academy of Sciences, 112(16), E2102–E2111.

Queller, D. C. (1997). Why do females care more than males? Proceedings of the Royal Society of London. Series B: Biological Sciences, 264(1388), 1555–1557.

Reynolds, J. D., Goodwin, N. B., & Freckleton, R. P. (2002). Evolutionary transitions in parental care and live bearing in vertebrates. Philosophical Transactions of the Royal Society of London. Series B: Biological Sciences, 357(1419), 269–281.

Schacht, R., Kramer, K. L., Székely, T., & Kappeler, P. M. (2017). Adult sex ratios and reproductive strategies: a critical re-examination of sex differences in human and animal societies. Philosophical Transactions of the Royal Society B: Biological Sciences, 372(1729), 20160309.

Simmons, L. W., & Kvarnemo, C. (2006). Costs of breeding and their effects on the direction of sexual selection. Proceedings of the Royal Society B: Biological Sciences, 273(1585), 465–470.

Stearns, S. C. (1976). Life-history tactics: a review of the ideas. The Quarterly Review of Biology, 51(1), 3– 47.

Stiver, K. A., & Alonzo, S. H. (2009). Parental and mating effort: Is there necessarily a trade-off? Ethology, 115(12), 1101–1126.

Storchová, L., & Hořák, D. (2018). Life-history characteristics of European birds. Global Ecology and Biogeography, 27(4), 400–406.

Székely, T., Webb, J. N., & Cuthill, I. C. (2000). Mating patterns, sexual selection and parental care: an integrative approach. In M. Apollonio, M. Festa-Bianchet & D. Mainardi (Eds.), Vertebrate mating systems (pp. 194–223). World Science Press.

Székely, T., Weissing, F. J., & Komdeur, J. (2014). Adult sex ratio variation: implications for breeding system evolution. Journal of Evolutionary Biology, 27(8), 1500–1512.

Trumbo, S. T. (2012). Patterns of parental care in invertebrates. In N. J. Royle, P. T. Smiseth & M. Kölliker (Eds.), The evolution of parental care (pp. 81–100). Oxford University Press.

Vági, B., Végvári, Z., Liker, A., Freckleton, R. P., & Székely, T. (2019). Parental care and the evolution of terrestriality in frogs. Proceedings of the Royal Society B, 286(1900), 20182737.

Yamamura, N., & Tsuji, N. (1993). Parental care as a game. Journal of Evolutionary Biology, 6(1), 103–127.

Zahavi, A. (1975). Mate selection—a selection for a handicap. Journal of Theoretical Biology, 53(1), 205–

