## Supplementary Information for "A life-history perspective on the evolutionary interplay of sex ratios and parental sex roles"

This Supplement includes four supplementary figures:

Figure S1. Sex differences in maturation time select for parental roles.

Figure S2. Overview of the effects of sex differences in maturation time.

Figure S3. Analytical predictions of the Fromhage-Jennions model.

Figure S4. Analytical predictions of the Fromhage-Jennions model (original parametrization).

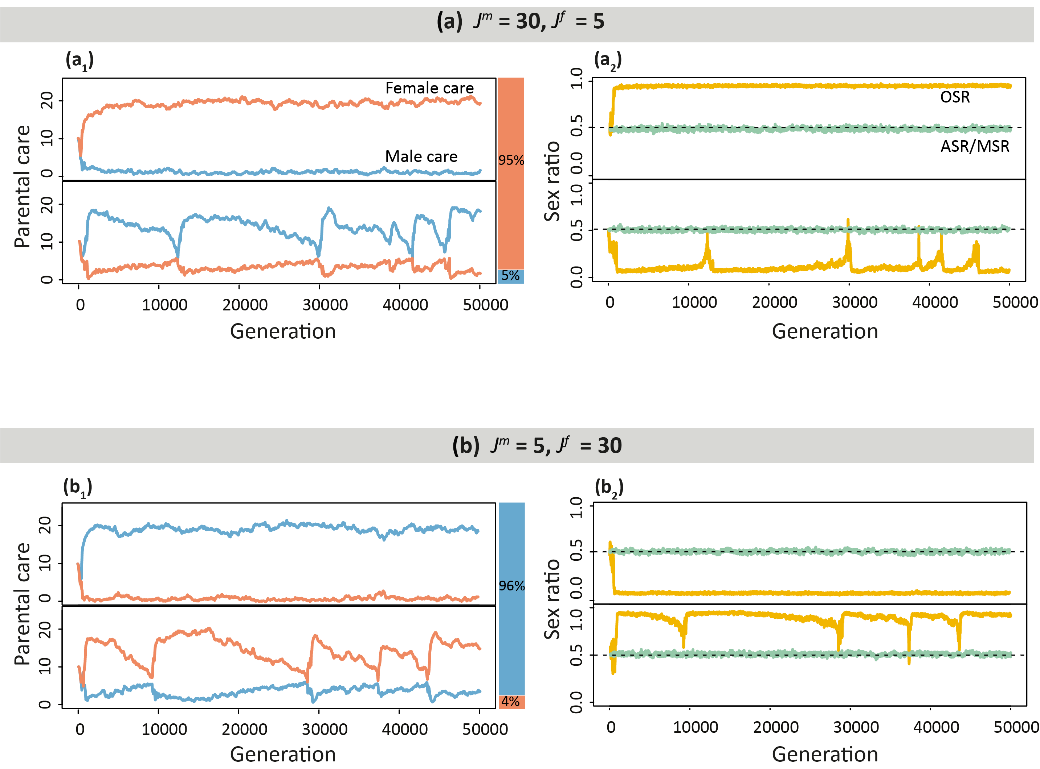

**Figure S1. Sex differences in maturation time select for parental sex roles.** The graphs depict a situation in which males and females mature at different rates (
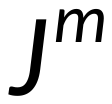
 and
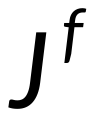
). Since all adult mortality rates are the same, the ASR is identical to the MSR. **(a)** When males mature relatively slowly (
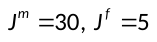
), female-biased care evolved in 95 of 100 simulations, together with a nearly unbiased ASR and a strongly male-biased OSR (the upper panels in (a_1_) and (a_2_) show a representative example). In 5 simulations (lower panels), male-biased care evolved, together with an almost unbiased ASR and a strongly female-biased OSR. **(b)** When females take a longer period to mature (
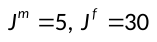
), male-biased care evolved in 96 of the 100 simulations (upper panels in (b_1_) and (b_2_)), together with a nearly unbiased ASR and a strongly female-biased OSR. In 4 simulations (lower panels), female-biased care evolved, together with an unbiased ASR and strongly male-biased OSR. Mortality rates are 0.001 for both sexes in all stages, and 100 replicate simulations with 50,000 generations were run per parameter setting. In all simulations parents had an additive effect on offspring survival (no synergy,
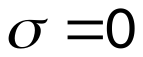
).

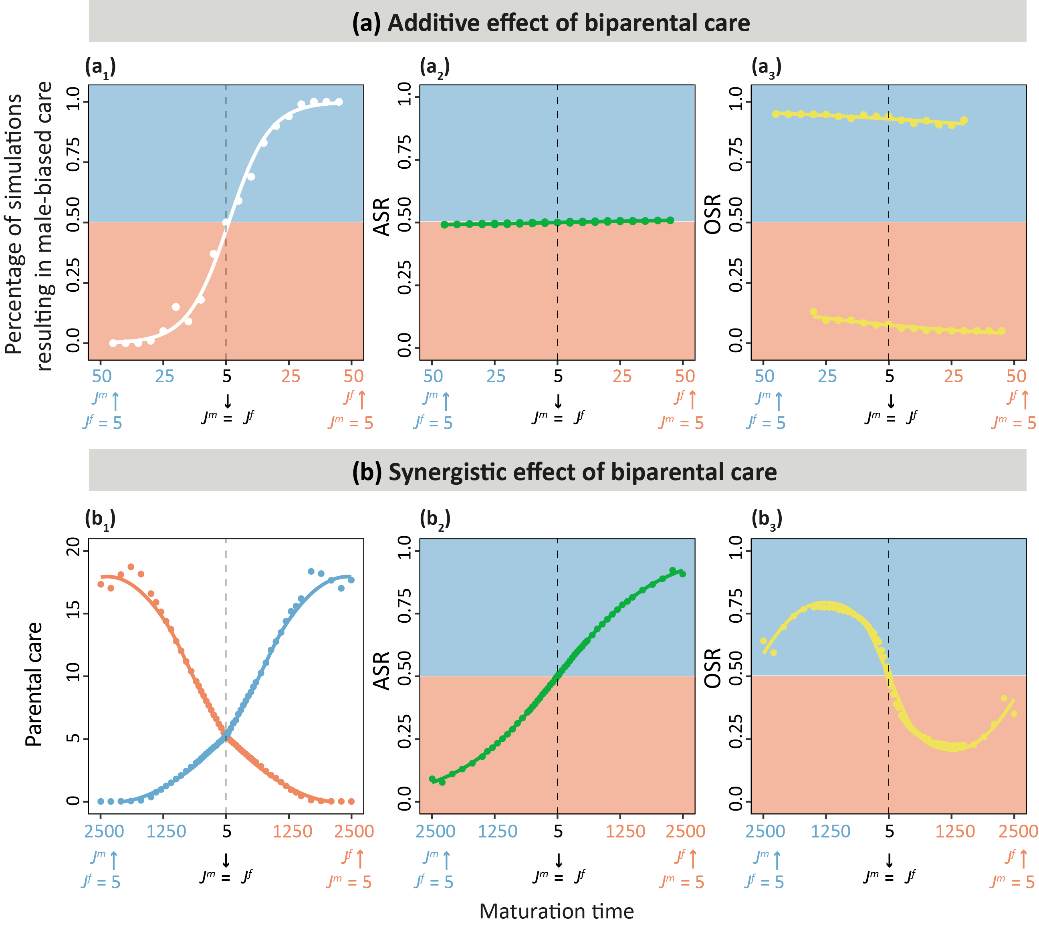

**Figure S2**. **Overview of the effects of sex differences in maturation time.** Outcome of a large number of simulations considering sex differences in maturation time. **(a)** In the absence of parental synergy (
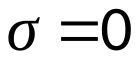
), when parents have an additive affect of offspring survival, all simulations resulted in either strongly male-biased or strongly female-biased care. The percentage of simulations resulting in either outcome depends on the direction and degree of the asymmetry in maturation times as shown in (a_1_): when males mature slowly, female-biased care is more likely to evolve, while male-biased care occurs more often when female juveniles spend a longer period to mature. The panel (a_2_) shows ASR is unbiased, unless the asymmetry in maturation times is very strong. The OSR is shown separately for the simulations resulting in male- and female-biased care in the panel (a_3_), in all cases, the OSR is skewed towards the less-caring sex. **(b)** In the presence of parental synergy (
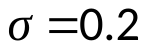
), biparental care evolves when sex-specific mortalities are not too different. Therefore, the panel (b_1_) now shows the average level of male and female care (averaged over 100 replicates). The sex that maturates faster tends to provide more care, and uniparental care evolves if the maturation time of the sexes is very different. In case of an extreme (more than 10-fold) asymmetry in maturation rates, the sex that maturates at a very slow pace has a larger chance of dying before reaching adulthood, resulting in an ASR bias in favour of the faster maturating sex. The OSR is biased toward the less-caring sex, and the relationship between the OSR and the degree of sexual asymmetry in maturation times is non-monotonic. 100 replicate simulations were run for (a) 50,000 generations and (b) 5,000 generation for each parameter setting. All simulations started from egalitarian care (
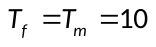
), each dot of the ASR, OSR (in a and b), male care level and female care level (in b) shown in the graph is the mean of 100 equilibrium outcomes. Notice that the horizontal scales of (a) and (b) are very different, we here show a considerably large range of asymmetry in maturation rate in (b) to make it comparable to Fig. 6.

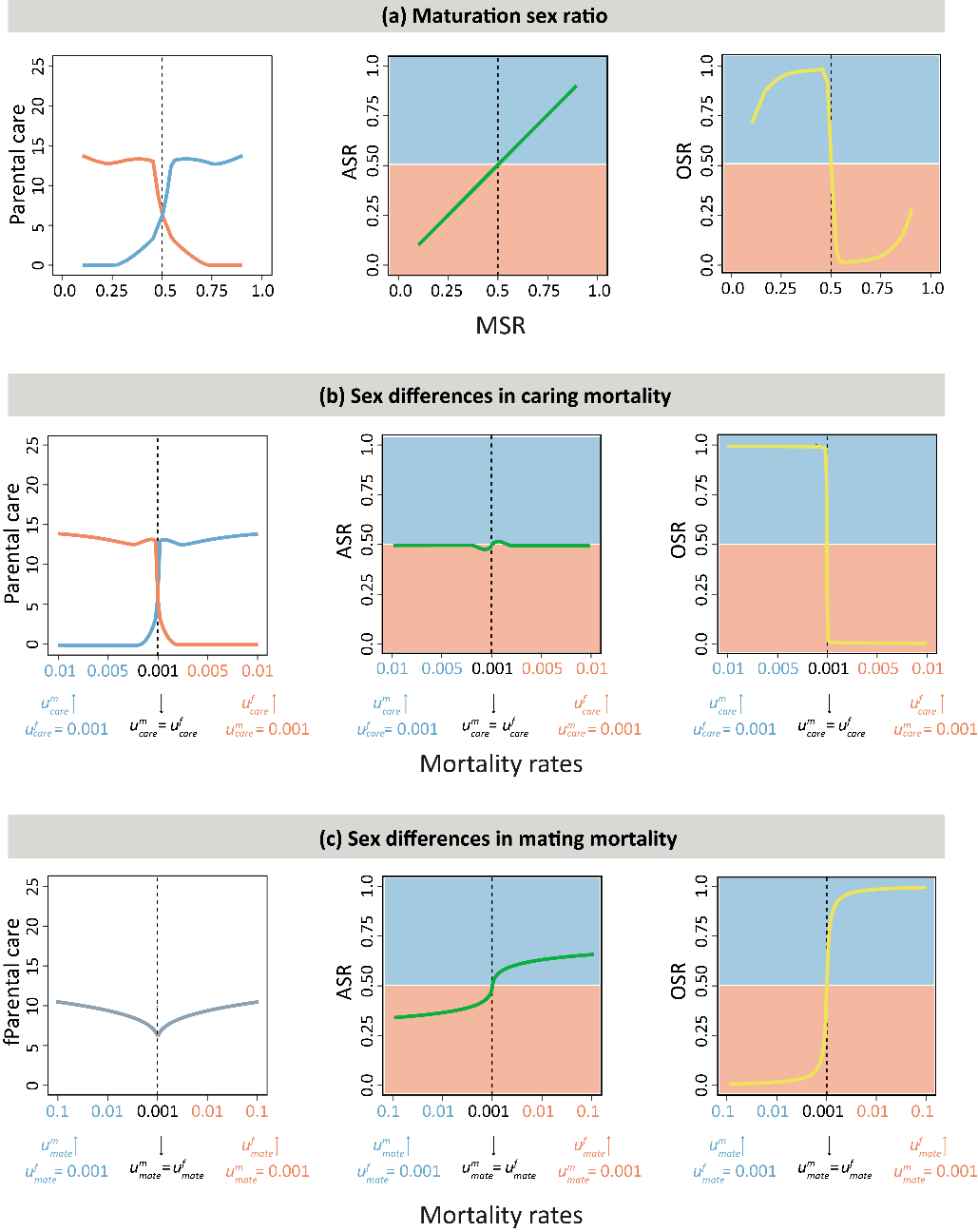
**Figure S3. Analytical predictions of the Fromhage-Jennions model.** By making use of the selection gradient method (evolution is supposed to proceed in the direction of steepest fitness ascent), Fromhage and Jennions (2016) were able to derive predictions on stable parental care levels and the corresponding ASR and OSR for various life-history scenarios. The analytical predictions could only be derived for the case of parental synergy. Although the Fromhage-Jennions model deviates from our model in several ways (see Long & Weissing, 2020 and below), all their conclusions concur with ours for the case of parental synergy (which are summarised in Fig. 6). **(a)** Fromhage and Jennions did not directly consider sex differences in juvenile mortality or maturation rate; instead, they used the maturation sex ratio (MSR), which is identical to the ASR in this scenario, as an indirect measure summarizing these differences. In line with Fig. 6(a) and Fig. S2(b), Fromhage and Jennions predict biparental care with a care bias toward the overrepresented sex under mild deviations from an even MSR, and uniparental care in case of large deviations. The predicted relationship between OSR and MSR also agrees well with the outcome of our simulations. When the sexes differ in **(b)** mating mortality rates or **(c)** caring mortality rates, the analytical predictions also agree very well with our simulations (see Fig. 6(b,c)). In the Fromhage-Jennions model, the offspring survival function is given by
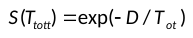
, where
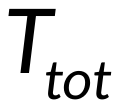
 is defined in a more complicated way than in our model:
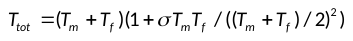
. Accordingly, the parental synergy parameter
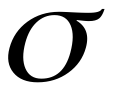
 has a slightly different meaning in our model. For the figure panels, we used the parametrization
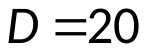
 and
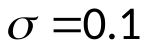
; moreover, all mortalities were, unless stated otherwise (along the horizontal axes), set to our default value 0.001. This choice aligns the two models relatively well (see Long & Weissing, 2020). For comparison, Figure S4 shows the predictions of the Fromhage-Jennions model for their original parametrization.

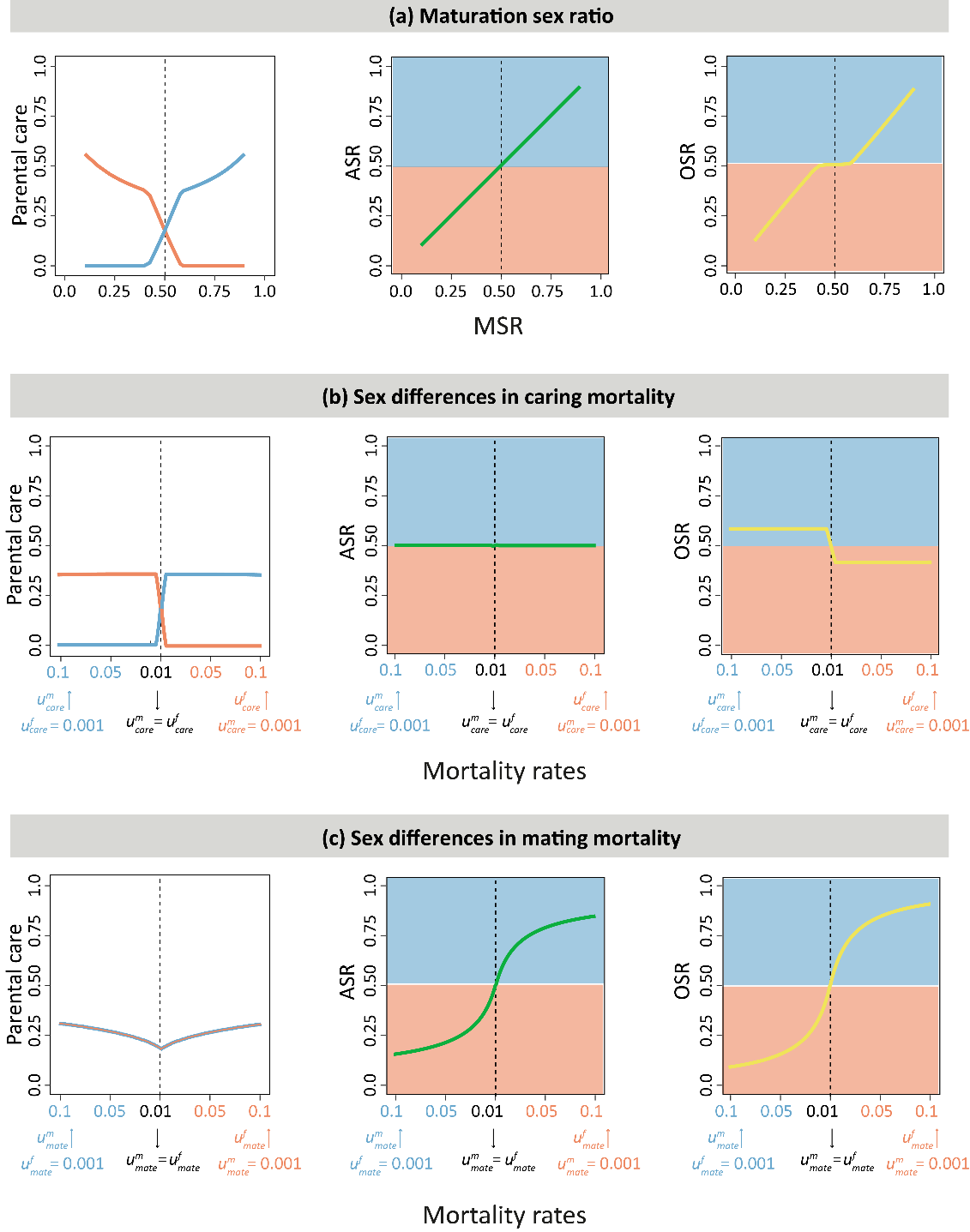
**Figure S4. Analytical predictions of the Fromhage-Jennions model (original parametrization).** Here we show the predictions of Fromhage and Jennions (2016) for the parametrization used in their article:
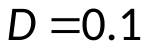
,
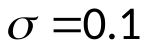
, and baseline mortality 0.01 (instead of our default value 0.001). Apart from some rescaling effects, the predictions concur well with those of Fig. S3, with two notable exceptions. First, the relationship between OSR and ASR (= MSR) is completely different in Figs. S3(a) and S4(a). This shows that same parental care pattern can be associated with very different OSR patterns. Second, the original parametrization predicts an even ASR in scenario (b) (sex differences in caring mortality), whereas Figs. S3(b) and Fig. 6(b) predict a small but systematic deviation from an even adult sex ratio. Fig. S4(b) shows that the evolution of pronounced parental sex roles is not necessarily associated with a bias in ASR (or MSR).
